# Familiarity modulates social approach toward stressed conspecifics in female rats

**DOI:** 10.1101/365312

**Authors:** Morgan M. Rogers-Carter, Anthony Djerdjaj, Amelia R. Culp, Joshua A. Elbaz, John P. Christianson

**Author notes:** Corresponding Author: Morgan M. Rogers-Carter.

## Abstract

Familiarity between conspecifics may influence how social affective cues shape social behaviors. In a social affective preference test, experimental rats, when given the choice to explore an unfamiliar stressed or a naive adult, will avoid interaction with a stressed conspecific. To determine if familiarity would influence social interactions with stressed conspecifics, male and female test rats underwent 2 social affective preference tests in isosexual triads where an experimental rat was presented with a naïve and a stressed target conspecific who were either familiar (cagemate) or unfamiliar. Male and female experimental rats avoided stressed unfamiliar conspecifics. However, experimental female rats demonstrated a preference to interact with their stressed, familiar cagemates. Male and female rats exhibited more self-grooming and immobility behavior in the presence of stressed conspecifics, which may indicate emotion contagion. These findings suggest a sex-specific role of familiarity in social approach and avoidance, and warrant further mechanistic exploration.

## Introduction

Animals can convey information about their emotional or physiological state via species-specific expressions, including vocalizations, chemosignals, olfactory cues and overt changes in behaviors. Thus, the generation of such cues by one individual and the subsequent detection and appraisal of these cues by observers enable the affect of one, or a few subjects, to influence the behavior of pairs or groups of animals [1]. The transmission of affect allows for the communication of impending threat between conspecifics [2] and exposure to a stressed animal alters the physiological state and behavior of an observer [3]. Observers may appraise situations to make decisions that can either protect them from harm, such as avoiding a sick or aggressive individual [4].

Identifying the mechanisms by which the social transmission of affect can alter the behavior of observers has received considerable attention. In a recent example, Sterley and colleagues report that when mice interacted with a recently stressed conspecific, the unstressed observer mice spent more time exploring their stressed counterpart, which was mediated by the paraventricular nucleus of the hypothalamus [5]. The ability to detect sickness, distress, or danger in another is evolutionarily adaptive because the observer may use this information to avoid illness or threat, which promotes survival and wellbeing. In rodents, both male and female mice avoid the bedding of male conspecifics with parasitic infections [6, 7] and male rats show decreased social exploration of both a male conspecific injected with lipopolysaccharide to mimic illness and avoid soiled bedding from similarly treated animals [8]. Male rats also avoid the soiled bedding of a footshocked conspecific [9] and mice will avoid the side of a chamber where odors from a footshocked conspecific were present [10].

The decision to approach or avoid another individual requires the integration of social emotional signals with situational and internal factors such as social rank, age and sex [11]. Familiarity between individuals is an important determinant in how a bystander responds to others in distress [3, 12, 13]. As an illustration, Signer and colleagues reported greater behavioral and neural responses in individuals viewing images of their loved ones in pain compared to similar images of strangers [14]. This powerful effect also relates to helping behaviors, and their neural correlates, which occur more frequently when help is directed toward in-group members than out-group members in distress [15]. In rodent paradigms in which an observer is confronted with a conspecific in pain or distress, familiarity is needed for, or augments, vicarious fear and empathic pain related behaviors [12, 16–22]. Regarding prosocial behavior, in the experiments reported by Burkett and colleagues, pair-bonded prairie voles demonstrated prosocial allogrooming of a stressed cagemate but not between unfamiliar voles [23]. Similarly, rats will learn to press a level to release a restrained conspecific if it belongs to a familiar group, but not if it is a different strain [24], but see [25]. In sum, the transmission, and perception, of distress between familiar individuals can cause a different pattern of behavior than transmission of the same distress to unfamiliar bystanders.

Improper detection of social communication or aberrant social decision-making may underlie the characteristic social impairments observed in numerous psychiatric orders. Schizophrenic participants exhibit impairments in theory of mind [26] and determination of trustworthiness [27]. Moreover, the inability to use social cues may correlate with the severity of social impairments in ADHD [28] and hinder one’s ability to detect how others respond to a socially undesirable behavior in autism [29]. While this is not a complete account of social decision making in psychiatric disorders [30] these examples underscore the need to understand
how social factors mediate social-decision making in order to develop more efficacious treatments for psychiatric conditions.

To investigate the role of factors like familiarity, sex and age in social responses to others in distress and the underlying neurobiology, we introduced a social affective preference (SAP) paradigm in which an observer rat is presented 2 conspecifics, 1 naive to treatment and the other stressed via footshock immediately prior to test. Experimental observer rats were free to interact with either of the targets. Consistent with other observations [8, 9] experimental adult rats avoided stressed conspecifics [31]. Notably, these observations were made in unfamiliar groups of rats. Here we used SAP tests to investigate the hypothesis that familiarity would promote social approach toward stressed conspecific rats. We conducted SAP tests in males and females with either unfamiliar or familiar (cagemate) interaction stimuli. Females approached stressed familiar conspecifics while all avoidance was predominant in all other treatment conditions.

## Methods

### Rats and Housing

36 female and 36 male adult (300g) Sprague-Dawley rats were obtained from Charles River Laboratories (Wilmington, MA), housed in isosexual groups of 4 with free access to food and water on a 12hr light/dark cycle. Experimental rats were randomly assigned to either a “Familiar” or “Unfamiliar” condition, resulting in 4 groups of experimental rats: females interacting with familiar conspecifics (Female-Cagemate), females with unfamiliar conspecifics (Female-Unfamiliar), males with familiar conspecifics (Male-Cagemate), and males with unfamiliar conspecifics (Male-Unfamiliar, n = 10 per group). The remaining 32 rats were designated as conspecific targets during social exploration testing. Experimental rats were housed and tested with isosexual conspecifics. Familiarity was established by cohousing experimental rats with conspecifics. Familiar group cages housed 2 experimental rats and 2 conspecifics; 1 of the conspecifics was used as the naive and the other as the stressed target for the SAP tests of the 2 experimental cagemates. Each conspecific cagemate pair was used for 2 SAP tests. The rats used for targets in SAP tests with unfamiliar conspecifics were housed separately in groups of 4. from their target conspecifics. All rats were housed in these assigned groups for 7 days prior to testing. Behavior tests were conducted within the first 4hr of the light phase and in accordance with the *Guide for the Care and Use of Laboratory Animals* and approved by the Boston College Institutional Animal Care and Use Committee.

### Social Affective Preference (SAP) Test

As described previously [31], the SAP test allows for the quantification of social exploration of an experimental rat directed toward 2 target conspecifics, 1 naïve to treatment and the other stressed. The SAP test began with 2 days of habituation: experimental rats were acclimated to the test area, a clear plastic cage (50 × 40 × 20cm, L × W × H) with a wire lid, for 60 min and then presented 2 empty restraint chambers (day 1) or 2 naive conspecifics (day 2) housed in the restraint chambers (see Fig. 1A). Restraint chambers were clear plastic enclosures (18 × 21 × 10 cm, L × W × H; see [32]) which allow for social exploration through acrylic rods spaced 1 cm center-to-center. To assess social affective preference on day 3 the experimental rat was presented 2 conspecifics, 1 naive to treatment and the other stressed via two footshocks (1mA, 5s duration, 60s inter-shock-interval; Precision Regulated Animal Shocker, Coulbourn Instruments, Whitehall, PA) immediately prior to testing in an adjacent room. SAP tests were recorded on digital video. Social exploration was defined as any time the experimental rat made physical contact with a conspecific and was quantified for both the naive and stressed targets. Social exploration was quantified during live testing and again by an observer who was completely blind to experimental conditions from digital video recordings to establish inter-rater-reliability (r_30_= 0.8528, p < 0.0001). From the digital video recordings, we constructed behavioral ethograms by quantifying of a number of experimental rat behaviors including: chamber exploration, bedding and perimeter investigation, escape-oriented behavior, immobility, self-grooming, and digging. Operational definitions for these behaviors are provided in Table 1. Chamber exploration, and investigative behaviors likely assess general locomotor function. Immobility is expressed in states of fear or anxiety [33], self-grooming may reflect emotion contagion [23] and biting and pulling may reflect aggression. The behavior of target rats was not quantified; interested readers are directed to our prior work for this analysis [31].

**Figure 1.**
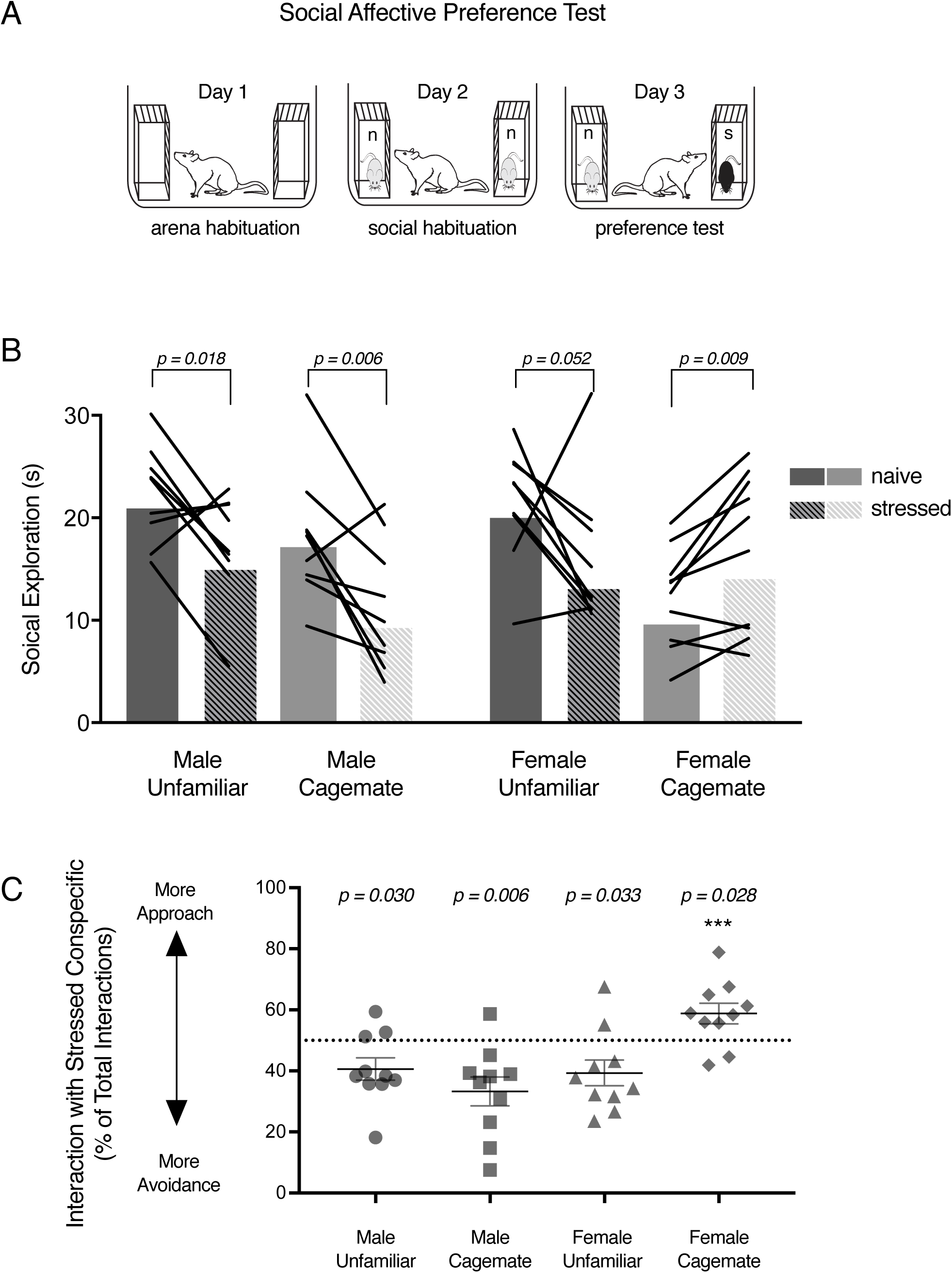
Familiarity mediates social avoidance of stressed conspecifics in female rats. (A) Schematic of the SAP test procedure with naive (n) conspecifics presented during social habituation and a naive and stressed (s) conspecific presented during SAP testing. Experimental rats received a series of 5-min exposures to the interaction arena over 3 days. (B) Mean (with individual replicates) social exploration time the experimental rats in each group engaged with naive or stressed conspecifics on Day 3. A sex-specific effect of familiarity was observed (F_1,36_ =8.479, p = 0.006): Rats in the Male-Unfamiliar (p = 0.018), Male-Cagemate (p = 0.006), and Female-Unfamiliar groups spent less time exploring the stressed conspecific compared to the naive (p = 0.052), whereas Female-Cagemate rats spent more time exploring the stressed conspecifics (p = 0.009). (C) Mean (individual replicates with ± s.e.m.) data from (B) expressed as the percent of social exploration time directed toward the stressed conspecific. Male-Unfamiliar (one-sample t_9_= 2.579, p = 0.029), Male-Cagemate (one-sample t,_9_= 3.550, p = 0.029), and Female-Unfamiliar (one-sample t_9_= 2.519, p = 0.032) percent preference scores were significantly less than 50%, indicating avoidance of the stressed conspecific. The Female-Cagemate percent preference score was greater than 50% (one-sample t_9_= 2.613, p = 0.028) indicating approach to the stressed conspecific; the Female-Cagemate preference score was significantly greater than the preference score of all other groups. ^∗∗∗^ p < 0.001.

**Table 1:**
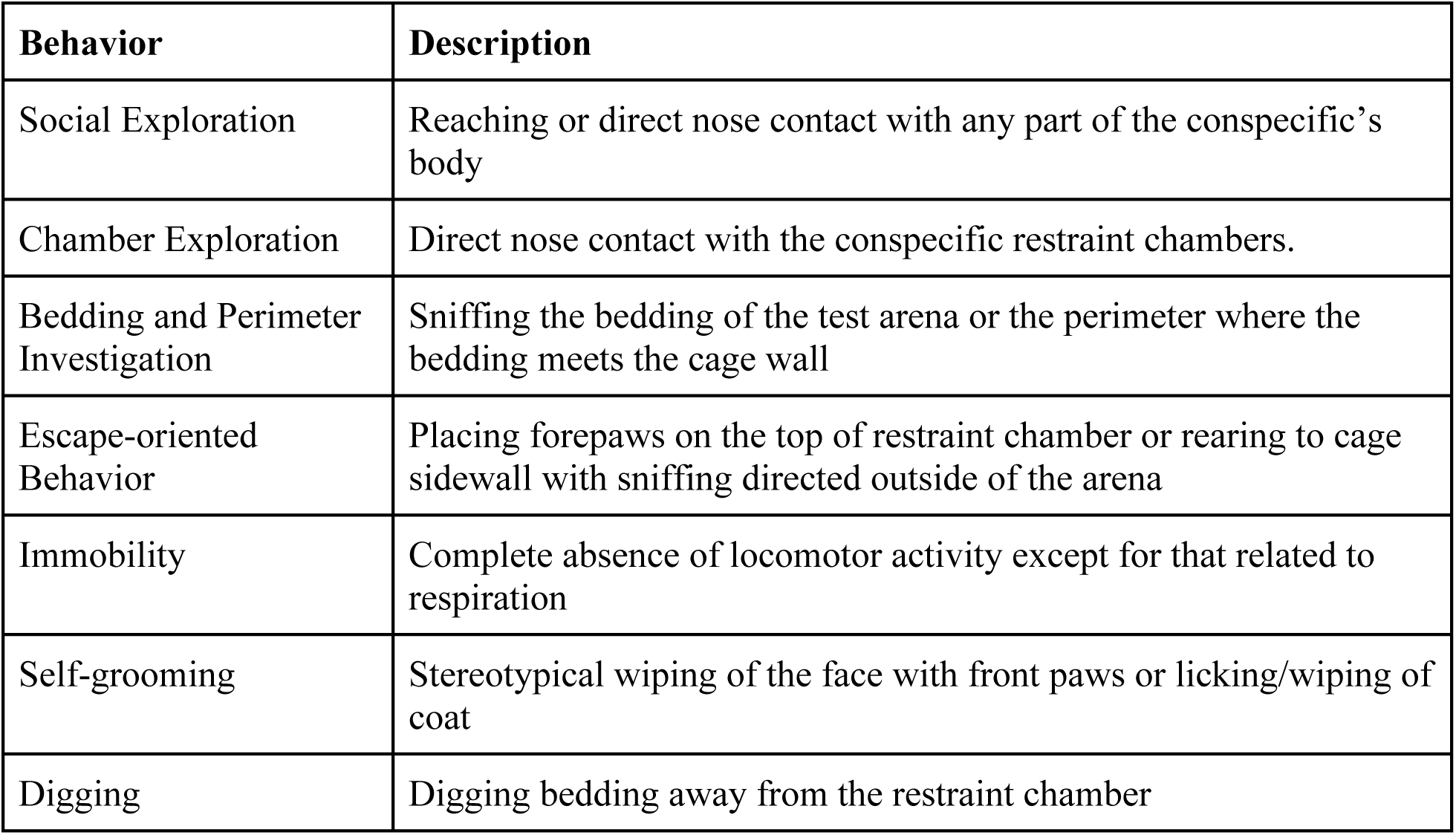
Description of behaviors quantified during SAP tests.

| Behavior | Description |
| --- | --- |
| Social Exploration | Reaching or direct nose contact with any part of the conspecific's body |
| Chamber Exploration | Direct nose contact with the conspecific restraint chambers. |
| Bedding and Perimeter Investigation | Sniffing the bedding of the test arena or the perimeter where the bedding meets the cage wall |
| Escape-oriented Behavior | Placing forepaws on the top of restraint chamber or rearing to cage sidewall with sniffing directed outside of the arena |
| Immobility | Complete absence of locomotor activity except for that related to respiration |
| Self-grooming | Stereotypical wiping of the face with front paws or licking/wiping of coat |
| Digging | Digging bedding away from the restraint chamber |

## Data Analysis

Experimental rat behaviors were analyzed with repeated measures 3-way ANOVAs in Prism 8.0 (GraphPad) with conspecific Affect (naïve or stressed) as a within-subjects factor, and Familiarity (familiar cagemates or unfamiliar conspecifics) and experimental rat Sex (male or female) as between-subjects factors. To account for individual differences in social exploration, a preference score was computed by dividing the time exploring the stressed conspecific by the total social exploration time (stress plus naive) times 100. Preference scores were compared to the hypothetical value of 50%, indicative of no preference, in a one-sample t-test and the effect of familiarity and experimenter sex on preference scores was tested with a 2-way ANOVA. Main effects and interactions were considered significant at p < 0.05 and experiment-wise type 1 error rate was maintained at α = 0.05 using Tukey or Sidak post hoc tests.

## Results

### Familiarity and social interaction with stressed conspecifics

Consistent with our previous findings [31] experimental rats spent less time exploring the unfamiliar stressed adult conspecifics compared to the naïve but female rats spent more time exploring familiar stressed conspecifics (Fig. 1b). This pattern resulted in a significant Affect X Familiarity X Sex interaction (F_1,36_ = 8.479, p = 0.006). Male experimental rats spent less time exploring the stressed conspecific compared to the naive for both unfamiliar (p = 0.018) and cagemate (p = 0.006) pairs. Female rats spent less time exploring unfamiliar stressed conspecifics (p = 0.052), but not stressed (p = 0.009) cagemates, when compared to naïve counterparts. To control for the range of individual variation in social exploration, we computed a preference score for the stressed conspecific (Fig. 1c). The preference scores were compared to 50%, the score at which there is no observable social affective preference. One-sample t-tests revealed preference scores significantly less than 50% for the Male Cagemate (t_9_ = 3.55, p = 0.006), Male Unfamiliar (t_9_ = 2.579, p = 0.029), and Female Unfamiliar (t_9_ = 2.519, p = 0.032) groups but significantly greater than 50% for the Female Cagemate (t_9_=2.613, p = 0.028) group. In sum, familiarity altered experimental female reactions to stressed conspecifics leading to greater social interaction.

### Analysis of experimental rat behavior

To explore if exposure to the stressed conspecific during SAP testing modulated experimental rat behavior, an observer blind to experimental conditions quantified behaviors displayed by the experimental rat on the social habituation and the SAP test sessions (Table 1). The proportion of time spent exhibiting each behavior was calculated by dividing the duration of time in a specific behavior by the total time of all behaviors (Fig. 2). 4 tests were not captured with digital video recording and therefore could not be analyzed resulting in the following sample sizes: Male-Unfamiliar n = 9; Male-Cagemate, n = 9; Female-Unfamiliar, n = 10; and Female-Cagemate, n = 8.

**Figure 2:**
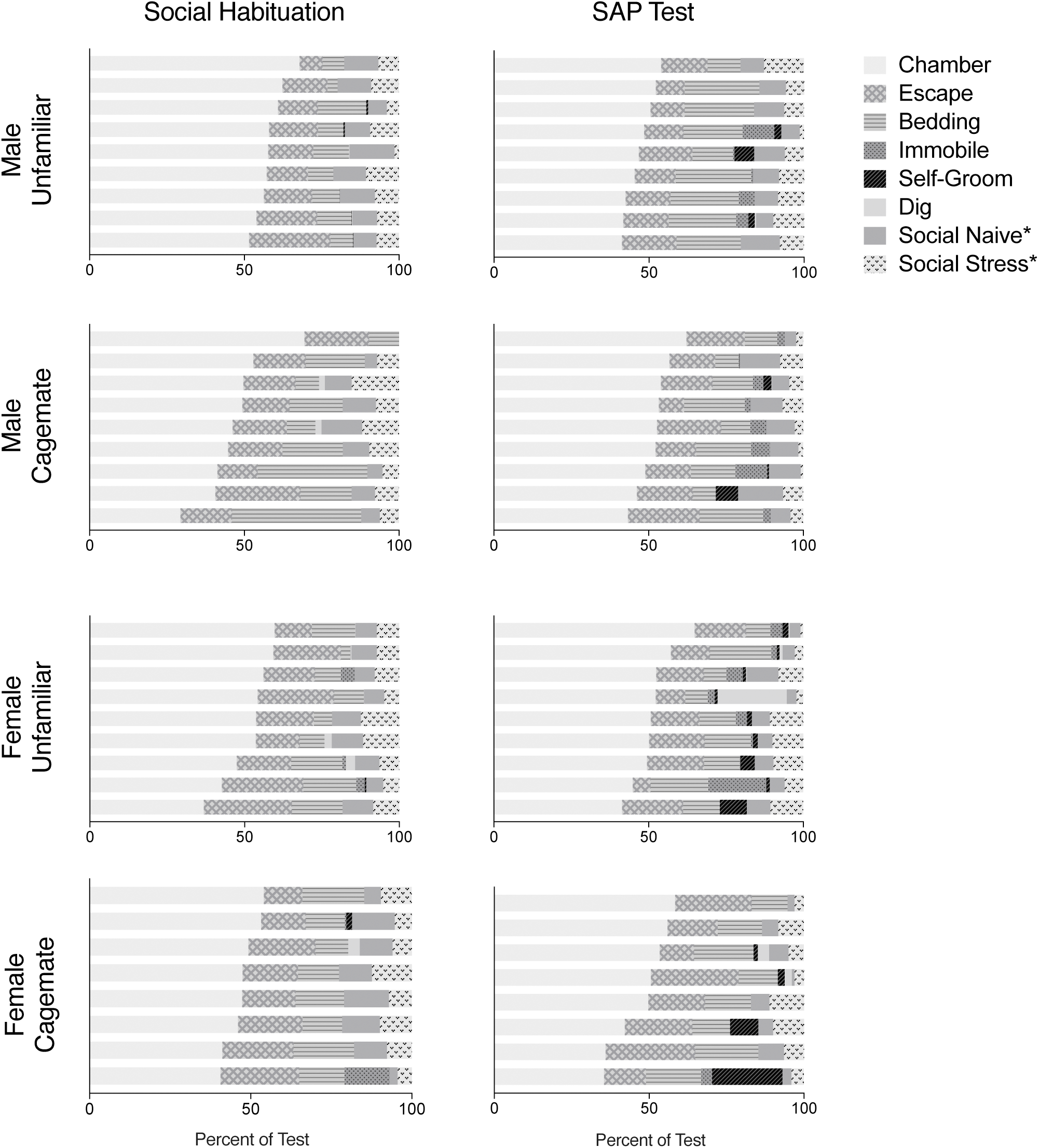
Behaviors observed during social habituation and SAP tests. Time spent engaged in chamber exploration, escape-oriented behaviors, bedding exploration, immobility, self-grooming, digging, and social exploration during the social habituation and SAP test days was quantified from digital video recordings (see Table 1). The sum of these behaviors was calculated for each rat and the data are plotted as the proportion of time each experimental rat demonstrated the behavior relative to the total. Each horizontal bar represents an individual subject. ^∗^Social exploration on habituation test day was directed at a naive conspecifics on the left or right side of the chamber.

3-way ANOVAs were performed for each quantified behavior. A significant Test Day X Familiarity X Sex interaction (F_1,56_ = 5.522, p = 0.022) and post-hoc comparisons on chamber exploration revealed that Male-Unfamiliar rats engaged in greater conspecific chamber exploration during habituation than any other group on either test day (Fig. 3a) which may reflect more social interest in this group compared to others. A main effect of Test Day (F_1,56_ = 9.404, p = 0.003) was found for self-grooming which reflected an increase in self-grooming during SAP testing than the social habituation test day in all treatment conditions (Fig. 3b). Similarly, a main effect of Test Day (F_1,56_ = 4.941, p = 0.030) and a Test Day x Sex interaction (F_1,56_ = 5.454, p = 0.231) on immobility indicated that experimental rats exhibit more immobility in the presence of a stressed conspecific than in the presence of naive conspecifics (Fig. 3c). For bedding and perimeter sniffing, the test revealed a main effect of Familiarity (F_1,16_ = 19.74, p < 0.001) indicating that experimental rats spent more time exhibiting non-social behaviors in the presence of cagemates regardless of sex or test day (Fig. 3d).

**Figure 3:**
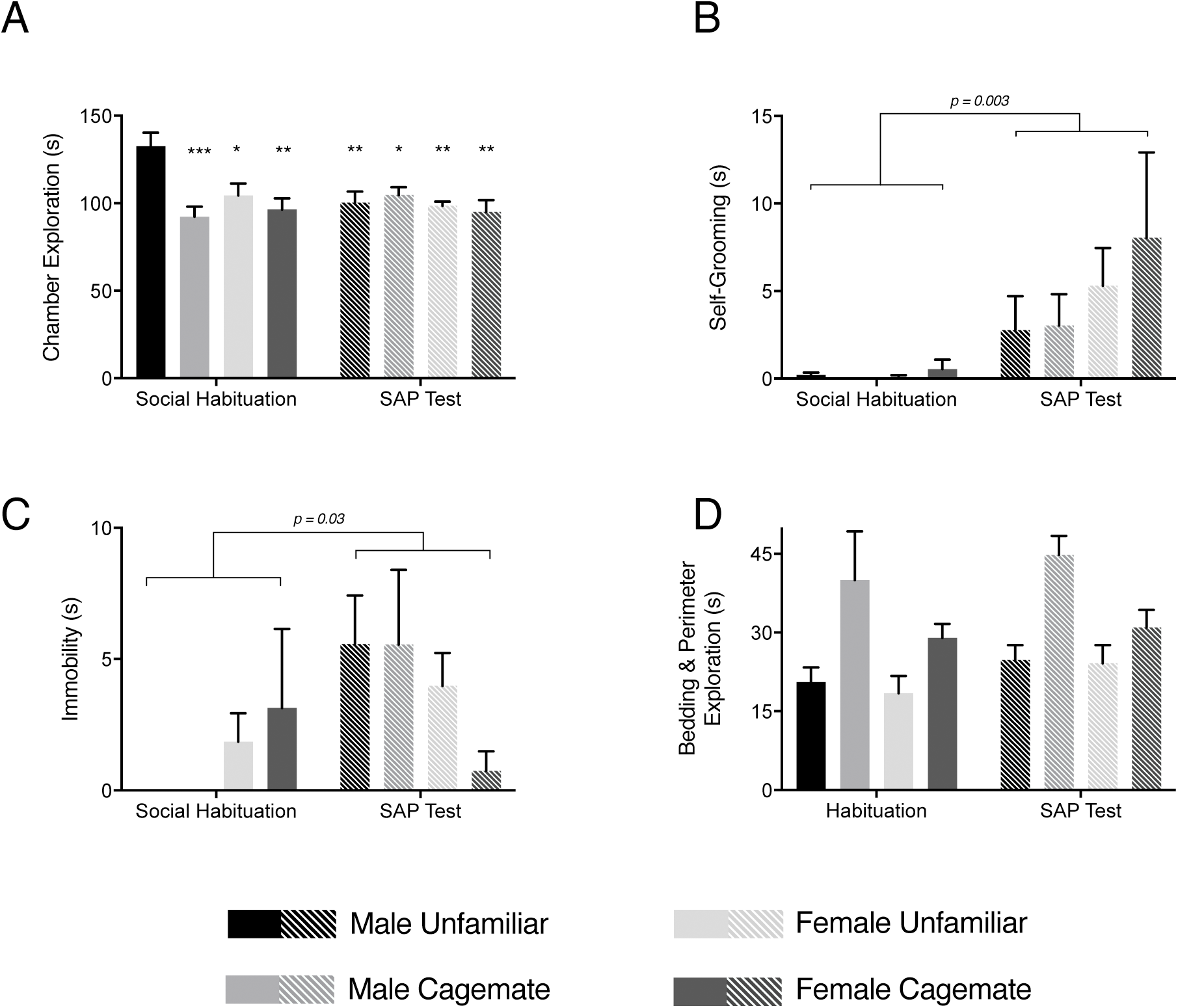
Comparison of experimental rat behaviors between habituation and SAP test days. (A) Mean (± s.e.m) time spent exploring the conspecific chambers on social habituation and SAP test days. Greater time spent in chamber exploration was observed in the Male-Unfamiliar group during habituation than any other group (F_1,56_ = 5.522, p = 0.022). ^∗^p < 0.05, ^∗∗^p < 0.01, ^∗∗∗^p<0.001 compared to Male-Unfamiliar. (B) Mean (± s.e.m) time spent self-grooming. In the presence of a stressed conspecific during SAP tests, experimental rats exhibited more self-grooming than during social habituation (F_1,56_ = 9.404, p = 0.003, main effect of Test Day). (C) Mean (± s.e.m) time experimental rats spent immobile, which was greater during SAP tests than social habituation (F_1,56_ = 4.941, p = 0.030, main effect of day). (D) Mean (± s.e.m) time experimental rats spent exploring the bedding or perimeter of the cage. Experimental rats spent more time exploring the bedding and perimeter during interactions with cagemates regardless of sex or day (F_1,16_ = 19.74, p = 0.000, main effect of Familiarity).

## Discussion

To explore if familiarity modulates social avoidance of stressed adult conspecifics, we compared adult male and female experimental rat social affective preference behavior when presented familiar versus unfamiliar adult conspecifics. Consistent with our previous findings, Male-Unfamiliar and Female-Unfamiliar experimental rats avoided stressed conspecifics [31]. Interestingly, we observed a sex-specific effect of familiarity. Female-Cagemate rats will approach stressed conspecifics, whereas Male-Cagemate rats still avoid the stressed conspecific. To our knowledge, this is the first evidence of sex differences regarding the role of familiarity on rat social emotional behaviors and suggests that males and females may appraise social stress signals in fundamentally different ways. That familiarity mediated social approach demonstrates that social preference behavior is likely not an innate, stereotyped response to conspecific affective cues, but rather a product of social decision making where affective cues are contextualized with factors like familiarity to inform situationally appropriate behavioral responses in sex-specific ways.

Social signals of stress may alert observers to looming threat. In the SAP test, rats may avoid stressed conspecifics because they are perceived to be dangerous. Consistent with this notion, we observed more frequent 22kHz ultrasonic vocalizations, which are thought to convey danger [1] during interactions between experimental adults and stressed adults than between experimental naive adults [31]. We also found that behaviors associated with states of fear and emotion contagion, immobility and self-grooming, were evident in SAP tests with males and with unfamiliar females in which we observed avoidance of the stressed conspecific (Fig. 3). Therefore, avoiding the stressed conspecific is likely a product of a conserved mechanism by which rodents use social communication to warn others of threat or harm.

That females will approach familiar, but not unfamiliar, stressed conspecifics suggests that familiarity modulates the way by which affective cues are integrated with other information during social decision-making in sex-specific ways. This pattern raises three questions regarding familiar females in the SAP test. First, do experimental females perceive or appraise the social stress signals of familiar rats differently than those of unfamiliar rats? Although insufficient data are available to completely address this question, female rats engaged in more rough and tumble play behavior with familiar than unfamiliar conspecifics but familiarity did not influence male behavior [34], consistent with a sex-specific effect of familiarity on the observer. Mikosz and colleagues reported increased amygdala and prefrontal cortex Fos expression in males and females in diestrus, but not females in estrus, after exposure to stressed conspecifics [35] suggesting a neural substrate for different perceptions of conspecific stress in males and females which, in turn, would provide a different neural milieu to integrate familiarity in females than in males. Second, do stressed rats emit different social signals to familiar observers than to unfamiliar? To our knowledge, this question has never been addressed but it is possible that the stress signals present in familiar female interactions, such as vocalizations or chemosignals, promote affiliative behaviors. In addition to the 22kHz danger signals, rodents emit a number of higher frequency vocalizations that are thought to facilitate social interaction, including maternal pup retrieval, and, in adults, reflect positive affective states [1]. Interestingly, Kiyokawa and colleagues isolated chemical compounds released by stressed rats that are sufficient promote either affiliation or avoidance [36]. Thus, it is possible that the combination of vocalizations, overt behaviors, and chemosignals generated by familiar female conspecifics is qualitatively different resulting in approach behavior.

Finally, does the presence of a familiar female observer buffer the stress of the demonstrator thus changing many aspects of the interaction? Demonstrators and observers have reciprocal effects on one another [5]. This idea is an important factor in regard to social buffering of fear, which is evident when the provision of a conspecific mitigates fear or stress. Social buffering occurs with several different stressors in many experimental contexts and has been observed in both male [37] and female [38] rats. Interestingly, social buffering is more robust between familiar rats [20] and animals of the same strain [39]. Although sex differences in social buffering have not been thoroughly explored [3, 40], it is possible that females are more effective than males at buffering fear in familiar conspecifics which might influence the reciprocal social behaviors and decision-making to give way to more social exploration.

Social behaviors are thought to be mediated by a decision-making process which requires the detection and evaluation of salient social stimuli to produce an appropriate social response. This adaptive process is mediated, in part, by the social decision-making network (SDMN; [41] which entails the social brain network [42] and the mesolimbic dopamine system [43]. The SDMN integrates environmental cues, internal physiological states, prior experience, and contextual and situational information to orchestrate specific social behaviors. The approach and avoidance behaviors that are sensitive to stress and familiarity reported here, therefore, are probably the product of neural activity in the SDMN. Following interaction with a stressed conspecific, increased Fos immunoreactivity is evident in a number of structures in the SDMN of observers including the amygdala [44], bed nucleus of the stria terminalis, and lateral hypothalamus [45]. Exposure to a stressed conspecific caused potentiation of glutamatergic synapses the paraventricular nucleus of the hypothalamus in observer rats suggesting that social stress signals change the excitability of SDMN circuits [5]. Moreover, the insular cortex and the anterior cingulate cortex, which are considered to be important to empathic processes in humans [46] and are anatomically connected with SDMN, have elevated Fos expression in observers and are required for social approach and avoidance responses to stressed conspecifics [23, 31]. Importantly, in autism spectrum disorders, abnormal resting state connectivity is observed in the amygdala and insula [47] and aberrant activation to social reward is documented in the anterior cingulate and mesolimbic structures [48]. Autism is characterized, in part, by impaired social function, and inappropriate social decisions may be the consequence of improper integration of situational factors and emotional states in SDMN structures. Thus, investigations into the behavioral and the neural mechanisms underlying social decisions will provide a basis for improving our understanding of the pathophysiology of aberrant social cognition and the advancement of more efficacious therapies.

## Acknowledgement

The authors have no competing financial or commercial interests to declare. Funding in support of this research was provided by the NSF Graduate Research Fellowship Program (M.M.R-C.); the Boston College Undergraduate Research Fellowships (A.C. & J. A.); the Gianinno Family (J.P.C) and NIH Grant MH109545 (J.P.C.). We would like to thank Nancy McGilloway and the Boston College Animal Care Facility Staff for the husbandry of research animals involved in this study.

